# Altered striatal long-term potentiation in the eIF4E- TG ASD mouse model

**DOI:** 10.64898/2026.03.04.709616

**Authors:** Alina Aaltonen, Jone Razquin Lizarraga, Julia Oyrer, Chiara Criscuolo, Ori Lieberman, Eric Klann, Anders Borgkvist, Emanuela Santini

## Abstract

Autism spectrum disorder (ASD) is associated with deficits in synaptic plasticity across brain regions. While striatal dysfunction is observed in various mouse models of ASD, the effect of ASD-associated genes on striatal plasticity has not been well characterised. We previously showed that overexpression of the SFARI ASD risk gene *eIF4E* in transgenic (eIF4E-TG) mice produces ASD-like behaviours and impairs dorsal striatal dopamine release. Here, we examined whether *eIF4E* overexpression alters striatal synaptic transmission and plasticity. Using microscopy, whole-cell electrophysiology, optogenetics and fast-scan cyclic voltammetry, we assessed dendritic morphology and excitatory synaptic properties of spiny projection neurons (SPNs). The eIF4E-TG mice exhibited higher dendritic spine density, elevated AMPA and NMDA receptor-mediated mEPSC frequency, and reduced AMPA mEPSC amplitude. We also observed an increased induction rate and magnitude of long-term potentiation (LTP) in SPNs, which is NMDA receptor-dependent but is not prevented by pharmacological D1 or D2 receptor antagonism under the conditions tested. Finally, we found that somatic and dendritic Ca^2+^ signals evoked by brief depolarisation are altered in SPNs from eIF4E-TG mice. Together, these findings are consistent with eIF4E overexpression promoting an NMDA receptor–dependent form of striatal LTP that is not prevented by D1/D2 receptor antagonism.

## Introduction

Synaptic plasticity occurs throughout the brain and underlies development and learning (Bliss and Lømo, 1973; Paolo Calabresi, Maj, Pisani, *et al*., 1992; Kirkwood, Rioult and Bear, 1996; Rogan, Staubli and LeDoux, 1997). The expression of synaptic plasticity in forms of long-term potentiation (LTP) and depression (LTD) is predominantly observed at the level of dendritic spines and synapses (Matsuzaki *et al*., 2004; Zhou, Homma and Poo, 2004; Harvey and Svoboda, 2007). LTP and LTD are regulated by many effectors and modulators, such as glutamatergic receptors (Collingridge, Kehl and McLennan, 1983; Oliet, Malenka and Nicoll, 1997) and neuromodulators (Burgard and Sarvey, 1990; Paolo Calabresi, Maj, Mercuri, *et al*., 1992; Calabresi *et al*., 1998; Cavaccini *et al*., 2018). In addition, the importance of protein synthesis in various aspects of synaptic plasticity has been established (Stanton and Sarvey, 1984; Frey *et al*., 1988; Kang and Schuman, 1996; Kelly, Mullany and Lynch, 2000; Bradshaw, Emptage and Bliss, 2003; Kelleher, Govindarajan and Tonegawa, 2004; Shrestha *et al*., 2020). Neuronal activity-dependent protein synthesis is thought to influence not only the magnitude of synaptic change but also the thresholds and persistence of plasticity, thereby shaping the rules by which synapses undergo long-term modification across circuits. These regulatory mechanisms operate across brain regions, but their functional consequences may differ depending on circuit architecture and neuromodulatory context.

Dysregulation of protein synthesis via mTORC1 signalling is implicated in ASD, both in idiopathic autism (Qin, Dai and Yin, 2016; Onore *et al*., 2017; Rosina *et al*., 2019) and associated genetic syndromes (Inoki *et al*., 2003; Butler *et al*., 2005; Hu *et al*., 2008; Kelleher and Bear, 2008; Sharma *et al*., 2010; Hoeffer *et al*., 2012; Longo *et al*., 2023). Mutations disrupting the mTORC1 pathway correlate with altered dendritic spine density and morphology across various brain regions (Hinton *et al*., 1991; Comery *et al*., 1997; Irwin *et al*., 2001; Nimchinsky, Oberlander and Svoboda, 2001; Galvez and Greenough, 2005; Koekkoek *et al*., 2005; Tavazoie *et al*., 2005; Grossman *et al*., 2006; Wang *et al*., 2022), as well as excitation-inhibition imbalance (Martin, Corbin and Huntsman, 2014; Braat *et al*., 2015) and numerous aberrant forms of synaptic plasticity (Huber *et al*., 2002; Li *et al*., 2002; Koekkoek *et al*., 2005; Zeng *et al*., 2007; Auerbach and Bear, 2010; Sharma *et al*., 2010; Chévere-Torres *et al*., 2012; Koga *et al*., 2015).

One circuit in which neuromodulatory context critically shapes plasticity is the striatum. Striatal dendritic spines, which harbour corticostriatal synapses, exhibit various forms of plasticity, including NMDA-receptor-dependent LTP (Spencer and Murphy, 2000; Perrin and Venance, 2019) and endocannabinoid (eCB)-dependent LTD (Gerdeman, Ronesi and Lovinger, 2002; Kreitzer and Malenka, 2005; Wu *et al*., 2015; Perrin and Venance, 2019). Many forms of striatal LTP/LTD are modulated by dopamine and show dopamine dependence that varies with induction protocol and circuit state (Paolo Calabresi, Maj, Mercuri, *et al*., 1992; Spencer and Murphy, 2000; Kreitzer and Malenka, 2005; Pawlak and Kerr, 2008; Reynolds *et al*., 2022). Studies in ASD models reveal disrupted corticostriatal functional connectivity (Shilyansky *et al*., 2010; Benthall, Ong and Bateup, 2018; Benthall *et al*., 2021; Pagani *et al*., 2021; Zerbi *et al*., 2021), suggesting that the striatum is affected by ASD-like synaptic plasticity dysregulation. In addition, Fragile X Syndrome (FXS) model mice have alterations in striatal spiny projection neurons (SPNs) including dendritic spines, altered actin turnover and striatal LTD, which correlate with repetitive behaviours and behavioural inflexibility (Jung *et al*., 2012; Longo *et al*., 2023; Mercaldo *et al*., 2023). TSC1 knockout also results in impaired eCB-LTD (Benthall *et al*., 2021). While there is a growing literature on striatal LTD in ASD models, it remains unclear whether long-term potentiation is similarly altered and whether such changes contribute to ASD-relevant circuit dysfunction. In particular, dysregulated striatal LTP is observed in mouse models of other striatal disorders, such as addiction disorders and Parkinson’s disease (Kitada *et al*., 2009; Ma *et al*., 2018). Given the central role of the striatum in action selection, habit formation, and behavioural flexibility, even subtle alterations in synaptic plasticity rules may have disproportionate consequences for circuit output and ASD-relevant behavioural phenotypes.

Eukaryotic initiation factor 4E (eIF4E) acts downstream of the mTORC1 pathway and is listed as a SFARI high-confidence ASD candidate gene (Simons Foundation, 2025), with rare variants implicated in autism (Neves-Pereira *et al*., 2009). eIF4E-TG (transgenic) and 4E-BP2 (repressor) knockout mice display ASD-relevant behavioural and synaptic phenotypes across brain regions (Gkogkas *et al*., 2013; Santini *et al*., 2013; Xu *et al*., 2020; Carbonell-Roig *et al*., 2024; Wang *et al*., 2024; Aaltonen *et al*., 2026). We previously identified enhanced striatal LTD in the eIF4E-TG mice (Santini *et al*., 2013), and more recently a deficit in dorsal striatal dopamine release (Carbonell-Roig *et al*., 2024), supporting altered striatal circuit function in this model. In the present study, we combined whole-cell patch clamp electrophysiology, optogenetics, fast-scan cyclic voltammetry and two-photon microscopy to test whether eIF4E overexpression alters excitatory synaptic structure and function in SPNs and modifies rules for long-term plasticity. We report increased SPN spine density, miniature current (mEPSC) frequency and reduced AMPA receptor-mediated mEPSC amplitude. While optogenetic stimulation of sensorimotor corticostriatal inputs did not reveal a change in baseline synaptic connectivity, we observed enhanced NMDA receptor–dependent potentiation induced by afferent stimulation under disinhibited recording conditions, which was not prevented by D1 or D2 receptor antagonism at the concentrations tested. Together, these data define an altered striatal plasticity profile in eIF4E-TG mice and motivate future work to determine the input specificity and neuromodulatory requirements of this potentiation.

## Results

### Increased dendritic spines in striatal SPNs of the eIF4E-TG mice

Altered dendritic spine density and morphology are reported in ASD postmortem studies (Hinton *et al*., 1991; Irwin *et al*., 2001). Consistently, dendritic spine alterations are also reported across multiple ASD mouse models and brain regions (Comery *et al*., 1997; Nimchinsky, Oberlander and Svoboda, 2001; Galvez and Greenough, 2005; Koekkoek *et al*., 2005; Tavazoie *et al*., 2005; Grossman *et al*., 2006; Peça *et al*., 2011; Wang *et al*., 2022; Longo *et al*., 2023). We have previously identified changes in dendritic spine density in other regions of the eIF4E-TG brain that correlated with altered synaptic function (Santini *et al*., 2013), while dendritic arborisation of SPNs was unchanged (Aaltonen *et al*., 2026). Because dendritic spine density in the striatum had not been assessed in this model, we quantified SPN spine density in the eIF4E-TG and WT mice. Individual SPNs were microinjected with 8 % Lucifer Yellow CH fluorescent dye and distal dendritic segments (50 µm from the soma) were analysed in 2-3-month-old male and female littermates. We observed a significant increase in spine density in SPNs of the eIF4E-TG mice (Fig. 1a-b). The increase in spine density was observed across dendrites, neurons and mice (Supplementary Fig. 1a) and is present in both D1- and D2-SPNs (Supplementary Fig. 1b-c).

**Figure 1.**
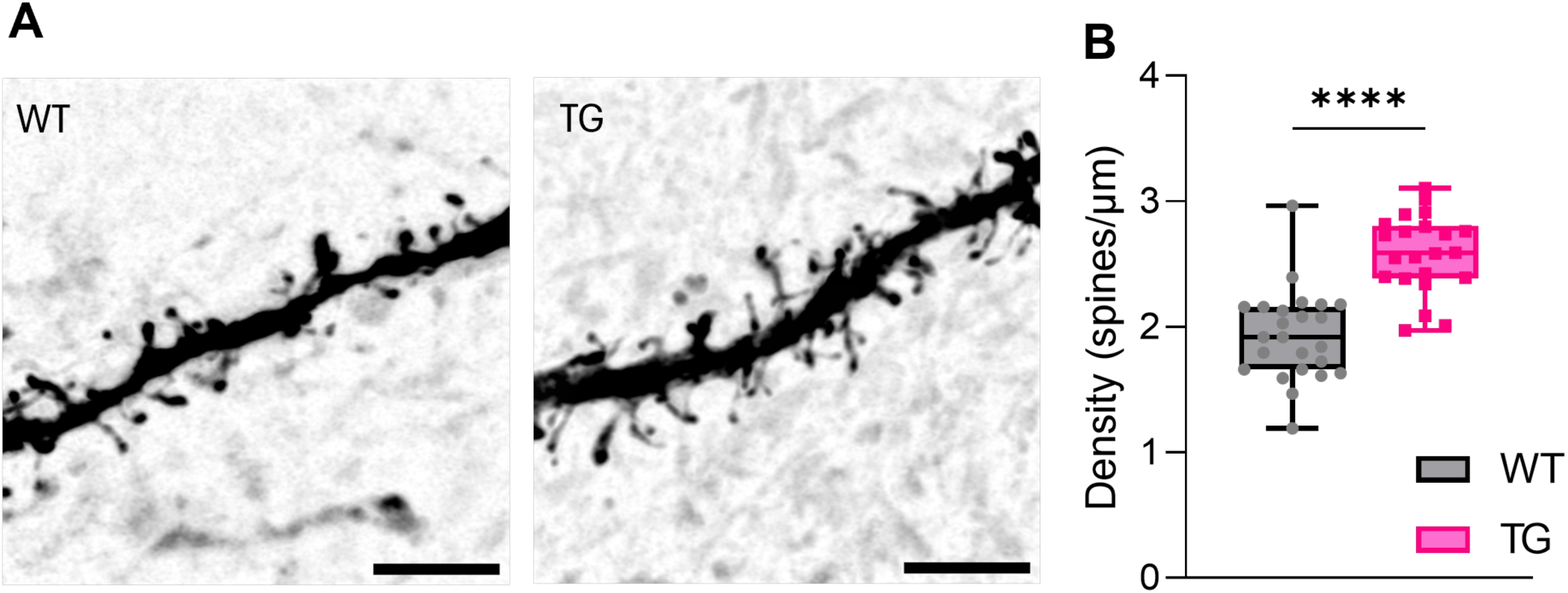
Dendritic spines of SPNs of eIF4E-TG mice. **A-B)** Images were collected from 3 WT and 3 TG mice (neurons n=24 WT, n=22 TG). **A)** Example Airyscan confocal images of SPN dendrites (scale bar 4 µm). **B)** SPN dendritic spine density in eIF4E-TG mice (unpaired T-test: t=6.56, ****p<0.0001). Data are presented as median ± quartiles within the box plot, with whiskers indicating minimum and maximum values. Dots represent individual values for each neuron. Significance is denoted as **** p<0.0001, calculated using Student’s t-test. For full details of statistical analysis, including negative results, refer to Supplemental table 1.

### Altered spontaneous synaptic transmission in the eIF4E-TG SPNs

As we observed an increase in dendritic spine density in the SPNs of TG mice, and distal dendrites on SPNs are preferentially innervated by excitatory afferents (Smith and Bolam, 1990; Huerta-Ocampo, Mena-Segovia and Bolam, 2014), we hypothesised that excitatory synaptic transmission may be altered. We performed voltage-clamp recordings in whole-cell configuration of miniature excitatory and inhibitory post-synaptic currents (mEPSCs and mIPSCs) in SPNs of 2-3-month-old D1-TdTomato/WT and D1-TdTomato/TG or D2-eGFP/WT and D2-eGFP/TG mice (see methods). As no differences were detected between D1- and D2-SPNs in dendritic spine density (Supplementary Fig. 1a-c) or in mEPSC/mIPSCs parameters (data not shown), recordings from both cell types were pooled for subsequent analyses.

To isolate AMPA receptor-mediated mEPSCs, recordings were performed at -70 mV in the presence of TTX to block sodium channels and prevent action potentials, and the GABAA-receptor antagonist picrotoxin to block inhibitory transmission. We found an increase in AMPA-mEPSC frequency and reduction in amplitude in eIF4E-TG mice (Fig. 2a-c), suggesting dysregulated spontaneous excitatory transmission.

**Figure 2.**
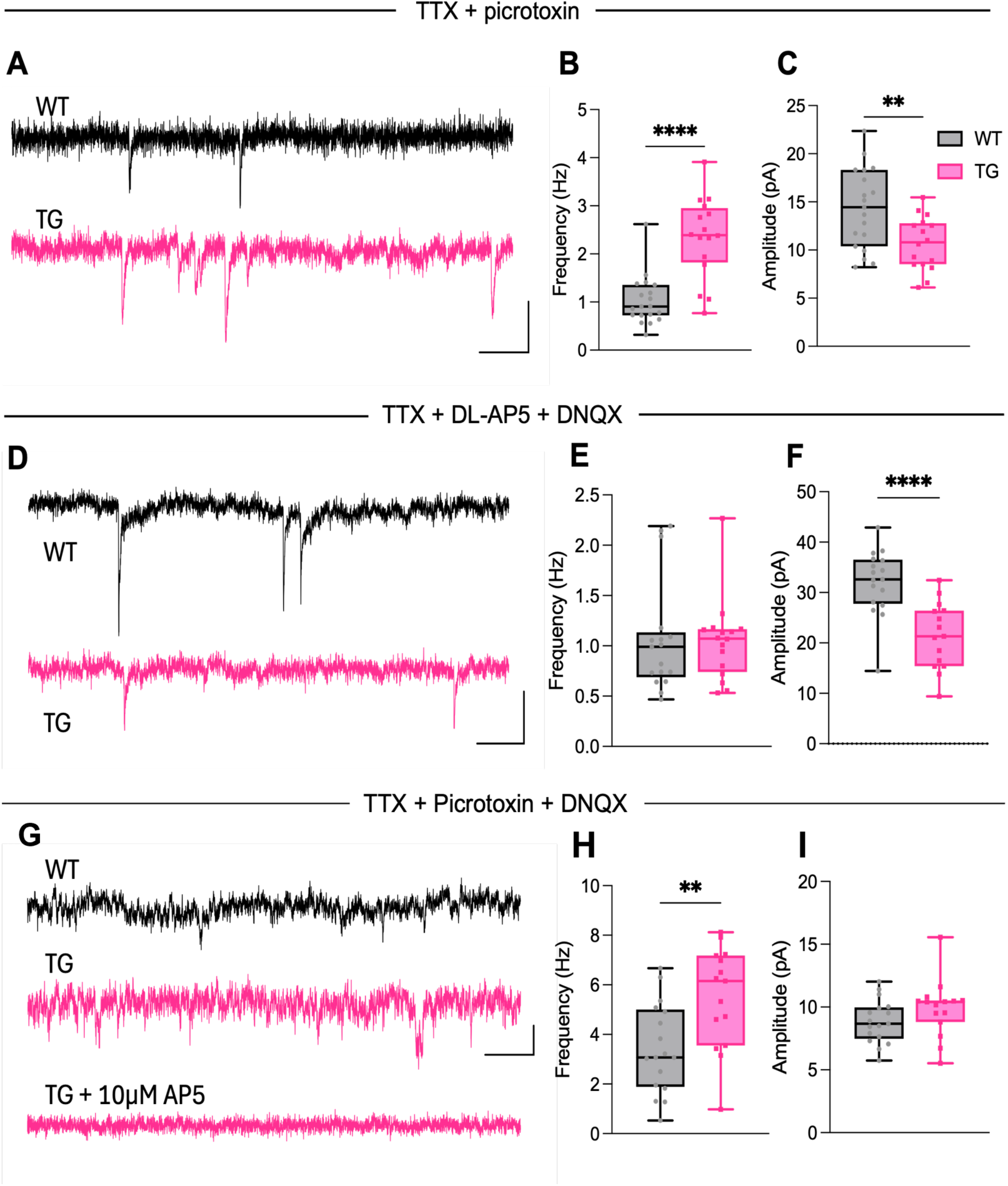
mEPSCs and mIPSCs from eIF4E-TG SPNs. **A-C)** Recordings were collected from 7 WT and 5 TG mice (neurons n=19 WT, n=16 TG). **A)** Representative mEPSC traces recorded with bath application of 1 µM TTX and 100 µM picrotoxin in intracellular solution (scale bar 20 pA, 200 ms). **B)** mEPSC frequency in eIF4E-TG mice (unpaired T-test: t=5.665, ****p<0.0001). **C)** mEPSC amplitude in eIF4E-TG mice (unpaired T-test: t=2.985, **p=0.0053). **D-F)** Recordings were collected from 8 WT and 5 TG mice (neurons n=17 WT, n=16 TG). **D)** Representative mIPSC traces recorded in presence of 1 µM TTX, 40 µM DL-AP5 and 10 µM DNQX (scale bar 20 pA, 200 ms). **E)** mIPSC frequency in eIF4E-TG mice. **F)** mIPSC amplitude in eIF4E-TG mice (unpaired T-test: t=4.587, ****p<0.0001). **G-I)** Recordings were collected from 5 WT and 6 TG mice (neurons n=17 WT, n=15 TG). **G)** Representative NMDA-mEPSC traces recorded in presence of 300 nM TTX, 50 µM picrotoxin and 10 µM DNQX. Bottom trace shows disappearance of events following bath application of 10 µM AP5 confirming NMDA mediation (scale bar 10 pA, 200 ms). **H)** NMDA-mEPSC frequency in eIF4E-TG mice (unpaired T-test: t=3, **p=0.0054). **I)** NMDA-mEPSC amplitude in eIF4E-TG mice. Data are presented as median ± quartiles within the box plot, with whiskers indicating minimum and maximum values. Dots represent individual neurons. Significance is denoted as ** p<0.01 and **** p<0.0001, calculated using Student’s t-test. For full details of statistical analysis, including negative results, refer to Supplemental table 2.

We next recorded mIPSCs in the presence of TTX, as well as AMPA-receptor and NMDA-receptor antagonists DNQX and AP5, using high-chloride internal solution to visualise inward GABAergic currents at -70 mV. We found that while the mIPSC amplitude was reduced in eIF4E-TG SPNs, mIPSC frequency was unchanged (Fig. 2d-f), suggesting an overall reduction in postsynaptic GABAergic signalling without changes in presynaptic inhibitory input. Taken together, these results indicate the presence of an excitatory-inhibitory imbalance in the inputs onto SPNs.

The increase in mEPSC frequency may reflect either an increase in presynaptic glutamate release or an increased number of excitatory synapses, whereas reduction in mEPSCs amplitude is consistent with altered postsynaptic AMPA receptor-mediated currents. As glutamatergic signalling is mediated by both AMPARs and NMDA-receptors (NMDARs), and it is possible to measure also spontaneous NMDAR-mediated transmission in SPNs (Vastagh *et al*., 2012), we then performed recordings in the presence of TTX, picrotoxin and AMPAR-antagonist DNQX and in Mg2+-free ACSF to unblock NMDARs at -70 mV. We found that the NMDA-mEPSC frequency was also enhanced in eIF4E-TG SPNs, while NMDA-mEPSC amplitude was unchanged (Fig. 2g-i). Together, these data demonstrate increased frequency of AMPAR- and NMDAR-mediated mEPSCs, as well as reduced amplitude of AMPAR-mediated mEPSCs and GABAR-mediated mIPSCs in SPNs of eIF4E-TG mice, indicating altered spontaneous excitatory and inhibitory synaptic transmission.

### Unaltered corticostriatal connectivity

Our findings of altered spontaneous excitatory transmission in SPNs raised the possibility that excitatory input to the striatum may be enhanced, either globally or in an input-specific manner. The striatum primarily receives excitatory synaptic input from the cortex (Ding, Peterson and Surmeier, 2008; Hunnicutt *et al*., 2016; Arias-García *et al*., 2018). Therefore, we sought to determine whether synaptic strength at defined corticostriatal inputs is altered in eIF4E-TG mice.

We targeted two cortical regions that provide among the highest proportions of excitatory input to the dorsal striatum: the primary motor cortex (M1) and primary somatosensory cortex (S1) (Wall *et al*., 2013). To selectively stimulate defined corticostriatal inputs, pAAV-hSyn-hChR2(H134R)-mCherry was bilaterally injected into the target cortical regions. Mice were allowed to recover for at least three weeks before preparation of acute striatal slices for whole-cell recordings. Photo-stimulation was then used to activate channelrhodopsin-expressing M1 or S1 axon terminals within the striatum. (Johansson and Silberberg, 2020).

Targeting specificity was assessed by examining mCherry expression in cortical regions using the Napari plug-in DMC-BrainMap (Jung *et al*., 2025). In M1-injected mice, 76.25 % of expression was localised to M1, with the remaining 23.75 % in M2 (Fig. 3a-b), indicating highly selective targeting of motor cortical regions. Optogenetic stimulation of M1 axon terminals reliably elicited optogenetic EPSCs (oEPSCs) in SPNs (Fig. 3c). When applying a set-intensity light stimulus, we observed no change in the oEPSC amplitude for motor-corticostriatal synapses (Fig. 3c). We also found no change in paired pulse ratio or NMDA:AMPA ratio of oEPSCs induced at M1-SPN synapses (Fig. 3d-f).

**Figure 3.**
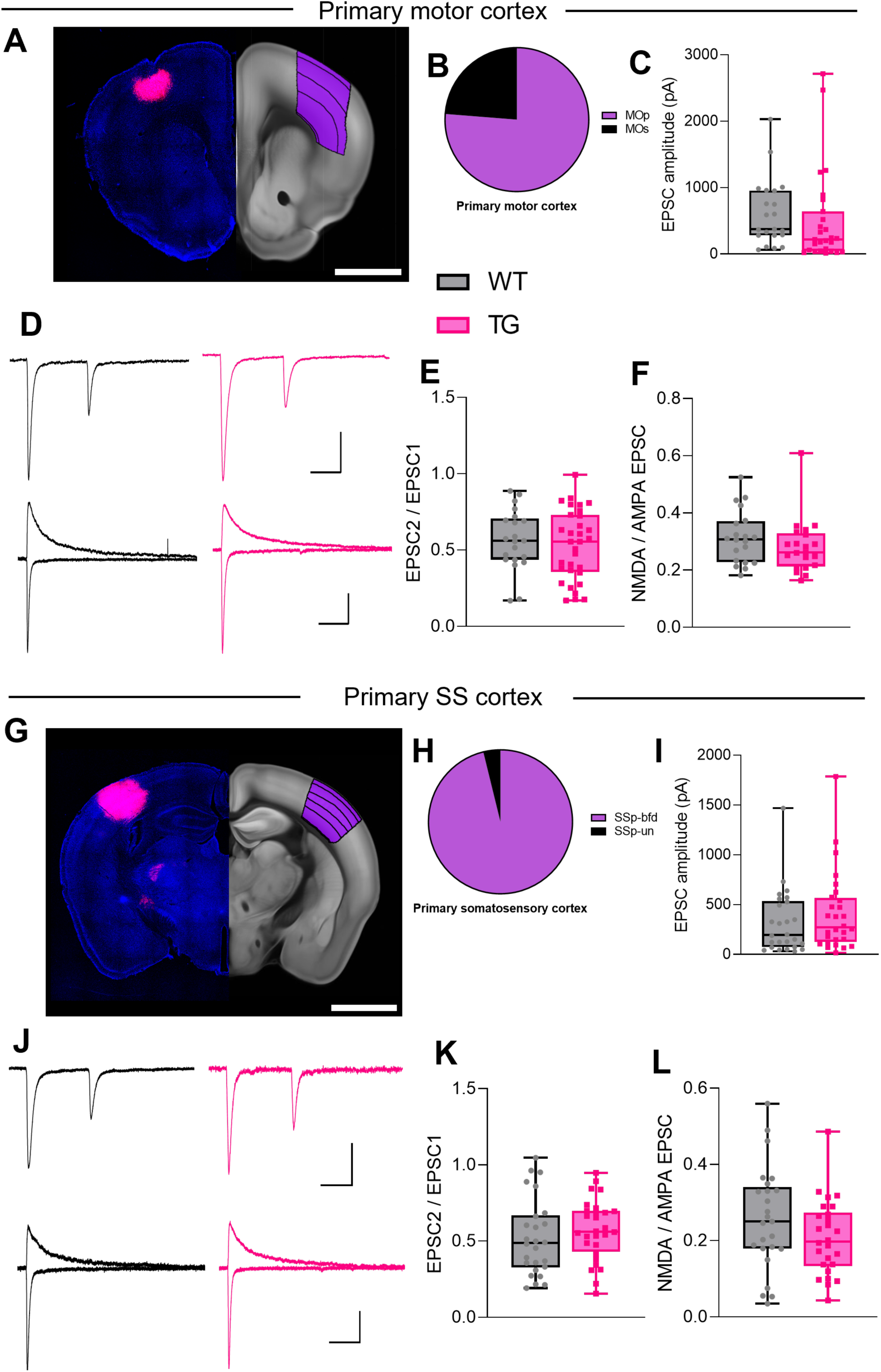
optogenetic stimulation of dorsal striatal inputs in eIF4E-TG mice. **A)** Left: example image of injection sites with mCherry expression (red) in a coronal brain slice at the level of the primary motor cortex, shown with DAPI neuronal staining (blue) (scale bar 2 mm). Right: equivalent coronal section from the Allen Brain Altas, with the target region indicated in purple. **B)** Pie chart depicting the averaged percentage of injection site localisation within the target region. MOp = primary motor cortex, MOs = secondary motor cortex. **C-F)** Recordings were collected from 4 WT and 7 TG mice (PPR: neurons n=21 WT, n=32 TG, NMDA: AMPA neurons n=20 WT, n=21 TG). **C)** Single pulse EPSC amplitude in eIF4E-TG mice with optogenetic stimulation of M1 axon terminals. **D)** Example traces showing paired-pulse EPSCs and NMDA: AMPA EPSCs from WT and TG mice (top: paired-pulse scale bars: 100 pA, 50 ms; bottom: NMDA: AMPA scale bars: 100 pA, 100 ms). **E)** Paired pulse ratio in eIF4E-TG mice. **F)** NMDA: AMPA ratio in eIF4E-TG mice. **G)** Left: example image of injection sites with mCherry expression (red) in a coronal brain slice at the level of the primary somatosensory cortex, shown with DAPI neuronal staining (blue) (scale bar 2 mm). Right: equivalent coronal section from the Allen Brain Altas, with the target region indicated in purple. **H)** Pie chart depicting the averaged percentage of injection site localisation within the target region. SSp-bfd = primary somatosensory area, barrel field, SSp-un = primary somatosensory area, unassigned. **I-L)** Recordings were collected from 9 WT and 7 TG mice (PPR: neurons n=26 WT, n=28 TG, NMDA: AMPA neurons n=25 WT, n=26 TG). **I)** Single pulse EPSC amplitude in eIF4E-TG mice with optogenetic stimulation of S1 axon terminals. **J)** Example traces showing paired-pulse EPSCs and NMDA: AMPA EPSCs from WT and TG mice (top: paired-pulse scale bars: 100 pA, 50 ms; bottom: NMDA: AMPA scale bars: 100 pA, 100 ms). **K)** Paired pulse ratio in eIF4E-TG mice. **L)** NMDA: AMPA ratio in eIF4E-TG mice. All recordings were performed in the presence of 50 µM picrotoxin to block GABAergic signalling. Statistical comparisons were performed using Student’s t-test. For full details of statistical analysis, including negative results, refer to Supplemental table 3.

Similarly, S1 injections resulted in 100 % localisation to primary somatosensory regions (Fig. 3g-h), again indicating highly selective targeting. We reliably elicited oEPSCs in SPNs with light stimulation of S1 terminals (Fig. 3i) but observed no change in the oEPSC amplitude in eIF4E-TG mice (Fig. 3i). We also found no change in paired pulse ratio or NMDA:AMPA ratio of oEPSCs at S1-SPN synapses (Fig. 3j-l). These findings indicate that glutamatergic synaptic strength and short-term presynaptic release properties at major sensorimotor corticostriatal inputs are preserved in eIF4E-TG mice under the conditions tested. Together, dendritic spine density measurements, spontaneous synaptic transmission recordings, and optogenetically assisted synaptic function assessments demonstrate that eIF4E overexpression increases excitatory synapses onto SPNs.

### Enhanced long-term potentiation

Previous studies in eIF4E-TG and 4E-BP2-knockout mice have reported altered forms of synaptic plasticity, including enhanced hippocampal LTP (Banko *et al*., 2005) and greater striatal LTD (Santini *et al*., 2013). In addition, we recently identified reduced striatal dopamine release in eIF4E-TG mice (Carbonell-Roig *et al*., 2024). Based on our observations of increased dendritic spine density, enhanced frequency of AMPAR- and NMDAR-mediated mEPSCs, and reduced AMPAR-mEPSC amplitude, we hypothesised that striatal SPNs in eIF4E-TG mice might display an altered plasticity profile. To test this, we performed whole-cell recordings with electrical stimulation of the white matter directly above the dorsal striatum (Fig. 4a). We used a high-frequency stimulation (HFS) protocol adapted from previous studies (Popescu, Saghyan and Paré, 2007; Wang *et al*., 2017; Zhou *et al*., 2019) consisting of four trains of 100Hz stimulation (1 s duration, 10 s interval), each paired with postsynaptic depolarisation to 0 mV (Fig. 4b).

**Figure 4.**
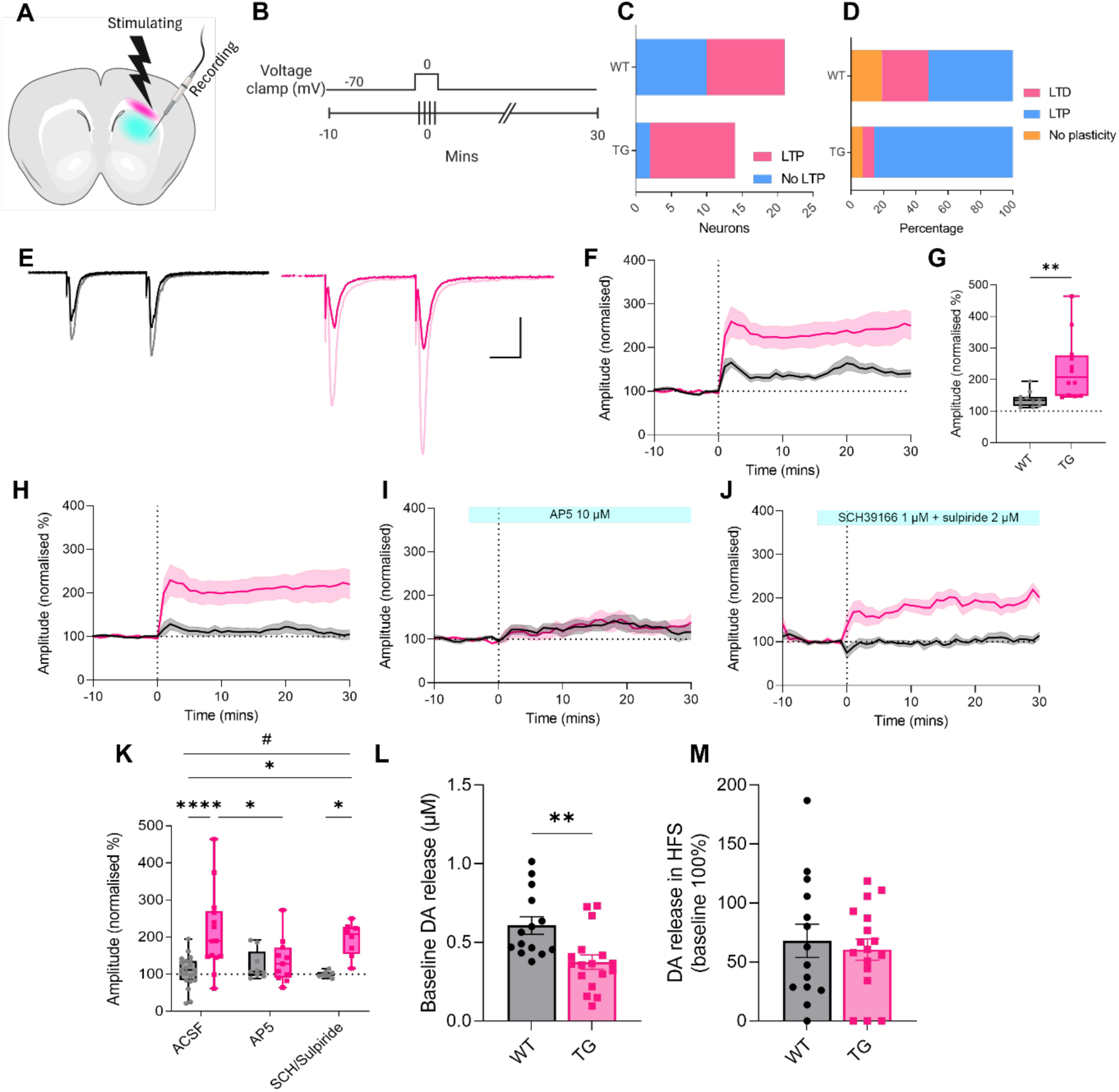
Plasticity induction with a high-frequency stimulation protocol in dorsal striatum of eIF4E-TG mice. **A)** Schematic showing the stimulating and recording positions in the dorsal striatum. Region used for stimulation (within corpus callosum) is depicted in magenta, and region for recording (within the dorsal striatum) is depicted in cyan. **B)** Schematic showing the stimulation protocol, with post-synaptic depolarisation paired with electrical high-frequency stimulation (HFS). Baseline was established over a 10-minute period prior to HFS, with paired EPSC (50ms inter-spike interval) delivered every 20 seconds, and EPSCs were recorded up to 30 minutes post-HFS. HFS was given at 100 Hz for one second, repeated four times with 10-second intervals. Voltage was clamped at -70 mV throughout recording but increased to 0 mV during each bout of HFS. **C-K)** In all recordings, baseline is normalised to 100 %, and plasticity was quantified from the final five minutes of recording (mins 25-30), compared to the final five minutes of baseline (mins -5-0). Plasticity was defined as LTD (<90 %), LTP (>110 %) or showing no change in EPSC amplitude from baseline (>90 % and <110 %). **C)** Proportion of cells exhibiting LTP (Chi-squared test: X^2^=4.413, *p=0.0418). **D)** Plasticity distribution: percentage of neurons expressing LTD, LTP or no change from baseline. **E)** Example traces showing paired-pulse EPSCs from LTP-expressing neurons from WT mice before (black) and after (grey) HFS, and from TG mice before (dark pink) and after (pale pink) HFS (scale bar: 200 pA, 20 ms). **F)** EPSC amplitude (normalised to baseline 100 %) for the duration of the recording period in cells expressing LTP (>110 % post-stimulation) from WT and TG mice. Recordings were collected from 10 WT and 12 TG mice (neurons n=11 WT, n=12 TG). **G)** EPSC amplitude 25-30 minutes after HFS (normalised to baseline 100 %) in cells expressing LTP (unpaired T-test: t=3.157, **p=0.0048). **H)** EPSC amplitude for the duration of the recording period in all cells from WT and TG mice in standard conditions (ACSF). ACSF recordings were collected from 21 WT and 13 TG mice (neurons n=21 WT, n=14 TG). **I)** EPSC amplitude for the duration of the recording period in all cells from WT and TG mice following bath application of 10 µM D-AP5. D-AP5 recordings were collected from 7 WT and 8 TG mice (neurons n=9 WT, n=11 TG). **J)** EPSC amplitude for the duration of the recording period in all cells from WT and TG mice following bath application of 1 µM SCH39166 and 2 µM sulpiride. SCH/sulpiride recordings were collected from 4 WT and 4 TG mice (neurons n=8 WT, n=8 TG). **K)** EPSC amplitude in all neurons from WT and TG mice, recorded in ACSF, D-AP5, or SCH39166/sulpiride (genotypeXdrug interaction, 2-way ANOVA: F(2, 65)=3.738, #p=0.0291). All recordings depicted in **A-K)** were performed in the presence of 50 µM picrotoxin to block GABAergic signalling. **L-M)** Fast-scan cyclic voltammetry recordings were obtained from 4 WT and 4 TG mice (slices n=14 WT, n=17 TG). **L)** Baseline dopamine (DA) release evoked with electrical stimulation (unpaired T-test: t=3.262, **p=0.0028). **M)** Dopamine (DA) release during HFS, expressed as a percentage of baseline release (100 %). Data are presented as median ± quartiles within the box plot, with whiskers indicating minimum and maximum values **(G,K)**, as mean ± SEM, with the mean represented by a connecting line and SEM by error lines with shading **(F,H-J)**, or as mean ± SEM, with bars indicating the mean and error bars depicting SEM **(L-M)**. Dots represent individual values for each neuron/slice. Significance is denoted as * p<0.05, ** p<0.01 and **** p<0.0001, calculated using Student’s t-test or Tukey’s post hoc multiple comparisons test, and # p<0.05, calculated using two-way ANOVA. The graphics in panels **D,F,H-J)** are included for illustrative purposes of the timeline of recording only, whereas statistical analysis is applied to the datasets in panels **C,G,K,L-M)**, as described. For full details of statistical analysis, including negative results, refer to Supplemental table 4.

In all experiments, LTP was defined as >10 % increase from baseline EPSC amplitude (>110 %) measured 30 minutes after stimulation, and LTD as >10 % reduction (<90 %). Our stimulation protocol induced LTP in both WT and TG mice, but we found an increased proportion of LTP-expressing neurons in the eIF4E-TG mice compared to WT (Fig. 4c). In WT mice, half of the recorded neurons exhibited LTP, with roughly a quarter showing LTD and a quarter showing no change from baseline, whereas over 80 % of neurons in TG mice expressed LTP (Fig. 4d). When analysing only LTP-expressing neurons, we found that eIF4E-TG SPNs exhibited greater post-stimulation EPSC amplitude (Fig. 4e-g). suggesting that in the TG mice, there is an increase in both the likelihood of LTP induction and in the magnitude of LTP. When all recorded neurons were included, WT neurons displayed bidirectional plasticity with no significant net change, whereas TG neurons showed a net potentiation (Fig. 4h, k), resulting in a significant genotype difference in post-stimulation EPSC amplitude (Fig. 4k). Together, these data indicate an increased likelihood and magnitude of potentiation in TG SPNs under this induction protocol.

As NMDARs are vital for the induction of many forms of LTP, including in the striatum (Collingridge, Kehl and McLennan, 1983; Spencer and Murphy, 2000; Perrin and Venance, 2019), we sought to determine whether this form of LTP also exhibits NMDAR dependency. The NMDAR antagonist AP5 (10 µM) was bath applied during baseline acquisition and HFS induction. Under these conditions, no difference in net plasticity was observed between WT or TG neurons (Fig. 4i, k), suggesting that LTP indication in this paradigm requires NMDAR activation.

Because dopamine signalling is critical to many forms of striatal plasticity (Paolo Calabresi, Maj, Mercuri, *et al*., 1992; Spencer and Murphy, 2000; Kreitzer and Malenka, 2005; Pawlak and Kerr, 2008; Reynolds *et al*., 2022), we next tested the effect of dopamine receptor antagonism by bath applying the D1R antagonist SCH39166 (1 µM) and the D2R antagonist sulpiride (2 µM). Under these conditions, TG neurons continued to exhibit net potentiation (Fig. 4j, k), whereas WT neurons did not express net plasticity, nor on a cellular level (Fig. 4k). Overall, these findings indicate that, LTP in eIF4E-TG SPNs was sensitive to NMDAR blockade but was not prevented by D1R/D2R antagonism.

As we previously observed reduced evoked dopamine release in the dorsal striatum of the TG mice (Carbonell-Roig *et al*., 2024), we asked whether altered dopamine dynamics during HFS could account for the persistence of LTP under dopamine receptor antagonism. Therefore, we used fast-scan cyclic voltammetry to record electrically evoked dopamine release in acute striatal slices during the HFS protocol described above. In line with our previous findings, we found a significant reduction in single pulse-evoked dopamine release (Fig. 4l). Surprisingly, dopamine release during HFS was detectable and did not differ from WT when expressed relative to baseline (Fig. 4m). This finding suggests that although baseline dopamine release is reduced in the eIF4E-TG mice, dopamine release during HFS is preserved and does not readily explain the continued expression of LTP in presence of dopamine receptor antagonists.

### Altered somatodendritic Ca^2+^ dynamics in eIF4E-TG SPNs

Synaptic plasticity in SPNs requires coordinated dopamine-dependent cAMP signalling and NMDAR-mediated Ca^2+^ influx (Carter and Sabatini, 2004; Shindou, Ochi-Shindou and Wickens, 2011; Jędrzejewska-Szmek *et al*., 2017). Because LTP in eIF4E-TG SPNs persisted under dopamine receptor antagonism, we asked whether depolarisation-evoked Ca²⁺ dynamics are altered in these neurons. We therefore used the genetically encoded Ca²⁺ indicator GCaMP7s and 2-photon (2P) microscopy to measure somatic and dendritic Ca²⁺ signals in SPNs from acute striatal slices following brief postsynaptic depolarisation.

pGP-AAV-syn-FLEX-jGCaMP7s-WPRE was bilaterally injected into the striatum of Adora2a-Cre/WT and -/eIF4E-TG or D1-Cre/WT and -/eIF4E-TG mice. After three weeks of recovery, acute striatal slices were prepared for imaging. SPNs were identified by baseline GCaMP7s fluorescence and voltage-clamped at -70 mV in whole-cell configuration. Cells were briefly depolarised to 0 mV in the absence of HFS to isolate postsynaptic Ca²⁺ responses.

Somatic and dendritic Ca²⁺ signals were imaged, with dendritic regions analysed 60 µm from the soma. Depolarisation-evoked GCaMP7s signals were reduced in both soma and dendrites of eIF4E-TG SPNs compared to WT (Fig. 5a-b). This effect was observed in both D1- and Adora2a-expressing cells (data not shown). Because absolute GCaMP fluorescence can vary across preparations, we additionally analysed dendritic responses relative to somatic responses within the same neuron. When expressed as a dendrite-to-soma ratio, Ca²⁺ signals were significantly increased in TG neurons (Fig. 5c), indicating an altered spatial distribution of depolarisation-evoked Ca²⁺ dynamics. Together, these findings demonstrate altered compartmental Ca²⁺ signalling in eIF4E-TG SPNs during depolarisation with a relatively larger Ca^2+^ influx in dendrites versus soma. Such changes in the spatial profile of Ca²⁺ responses may influence how synaptic inputs are integrated during plasticity-inducing conditions.

**Figure 5.**
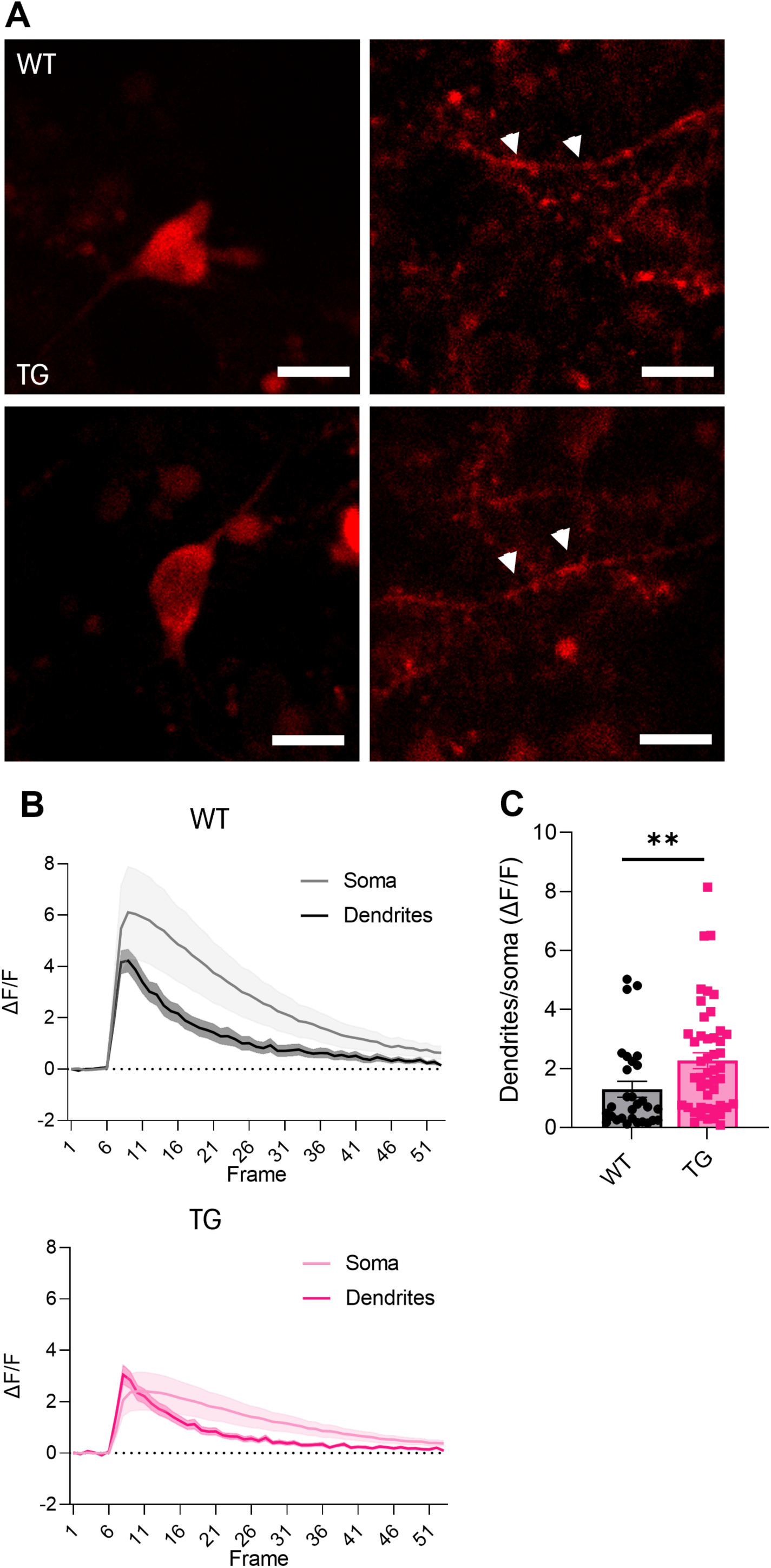
Calcium dynamics in SPNs of eIF4E-TG mice. **A)** Representative images of GCaMP7s activity in soma and in dendrites (60 µm away from soma) during depolarisation of SPNs (scale bar 10 µm). White arrows indicate dendritic signal observed during depolarisation. **B)** Time course of GCaMP7s activity in soma and dendrites for WT (left) and TG (right). Recordings were collected from 4 WT and 5 TG mice (neurons n=11 WT, n=16 TG; dendrites n=30 WT, n=47 TG). **C)** Ratio of dendritic to somatic GCaMP7s peak amplitude upon depolarisation (Mann-Whitney test: U=422, **p=0.0051). Data are presented as mean ± SEM, with the mean represented by a connecting line and SEM by error lines with shading **(B)**, or as mean ± SEM, with bars indicating the mean and error bars depicting SEM **(C)**. Significance is denoted as * p<0.05, ** p<0.01 and **** p<0.0001, calculated using the Mann-Whitney test. The graphics in panel **B)** are included for illustrative purposes of the timeline of recording only, whereas statistical analysis is applied to the datasets in panel **C)**, as described. For full details of statistical analysis, including negative results, refer to Supplemental table 5.

## Discussion

In this study, we identify an altered striatal synaptic phenotype in eIF4E-TG mice characterised by increased dendritic spine density, changes in spontaneous excitatory and inhibitory transmission, and enhanced NMDAR-dependent potentiation induced by high-frequency stimulation. Although baseline synaptic strength at major sensorimotor corticostriatal inputs was preserved, SPNs from TG mice displayed an increased likelihood and magnitude of LTP under disinhibited recording conditions. Notably, this potentiation was sensitive to NMDAR blockade but was not prevented by D1/D2 receptor antagonism at the concentrations tested. In parallel, we observed altered compartmental Ca²⁺ dynamics during postsynaptic depolarisation, suggesting that eIF4E overexpression modifies both structural and activity-dependent properties of striatal SPNs.

### Altered excitatory and inhibitory synaptic transmission

We observed an increase in dendritic spine density accompanied by an increase in mEPSC frequency, consistent with an increased number of excitatory synaptic contacts onto SPNs. The elevation in mEPSC frequency was detected for both AMPAR- and NMDAR-mediated events, indicating enhanced baseline glutamate signalling. Increased mEPSC frequency can reflect either higher presynaptic release probability or an increased number of functional synapses; however, the parallel increase in dendritic spine density supports the possibility of increased synapse number in eIF4E-TG SPNs. Similar increases in spine density and mEPSC frequency have been reported in other syndromic models of ASD characterised by elevated mTORC1-signalling and neuronal protein synthesis, including FXS mice, although in that case the effect was restricted to D1-SPNs (Longo *et al*., 2023). Increased mEPSC frequency has also been observed across multiple ASD models (Portmann *et al*., 2014; Peixoto *et al*., 2016), suggesting that elevated spontaneous excitatory transmission may represent a convergent phenotype.

In contrast, we observed a reduction in AMPA-mEPSC amplitude, which may indicate altered postsynaptic AMPAR function or composition and could reflect the presence of weaker or immature synapses. Consistent with this interpretation, FXS mice show dysregulated AMPAR GluR1/2 subunit expression (Muddashetty *et al*., 2007) and immature striatal dendritic spines morphology (Mercaldo *et al*., 2023), suggesting that increased synapse number may coexist with altered synaptic maturation in mTORC1-related ASD models.

We also identified a reduction in mIPSC amplitude without a change in mIPSC frequency in the eIF4E-TG mice. While this pattern is consistent with altered postsynaptic GABAergic signalling, our data do not provide direct evidence regarding changes in inhibitory synapse number or presynaptic release probability. Together, the combined alterations in spontaneous excitatory and inhibitory transmission are compatible with a shift in excitatory-inhibitory imbalance in the striatum of the eIF4E-TG mice. Similar alterations in striatal excitatory-inhibitory balance have been reported in various ASD models including FXS mice (Rothwell *et al*., 2014; Braat *et al*., 2015; Horder *et al*., 2018) as well as clinical studies (Horder *et al*., 2018). However, because corticostriatal and plasticity recordings were performed in the presence of picrotoxin, the functional contribution of altered inhibitory signalling to synaptic plasticity in the striatum of the eIF4E-TG mice remains to be determined. Future work should examine potential changes in striatonigral, striatopallidal, and intrastriatal inhibitory transmission.

### Preserved corticostriatal connectivity

Altered corticostriatal connectivity has been widely associated with various forms of ASD, both in clinical (Menon *et al*., 2004; Turner *et al*., 2006; Hoeft *et al*., 2007; Shilyansky *et al*., 2010; Di Martino *et al*., 2011; Abbott *et al*., 2018; Balsters, Mantini and Wenderoth, 2018; Pagani *et al*., 2021) and preclinical studies (Shilyansky *et al*., 2010; Kuo and Liu, 2017; Benthall, Ong and Bateup, 2018; Pagani *et al*., 2021; Zerbi *et al*., 2021; Longo *et al*., 2023). Besides the increase in spontaneous excitatory transmission, we did not detect changes in paired-pulse ratio or NMDA:AMPA ratio at motor or somatosensory corticostriatal inputs. These findings indicate that major sensorimotor corticostriatal synapses exhibit preserved baseline transmission in eIF4E-TG mice. However, our experiments were restricted to defined sensorimotor inputs, and other cortical or subcortical afferents to the striatum were not examined. In addition, an increase in the number of synaptic contacts would not necessarily be reflected in measures of synaptic strength during evoked stimuli, which produces synchronised glutamate release from multiple synapses. Our results suggest that enhanced spine density and mEPSC frequency may coexist with preserved evoked synaptic strength at specific input pathways.

### Dysregulated plasticity in the striatum

SPNs express bidirectional plasticity in both D1- and D2-SPNs (Akopian *et al*., 2000; Fino, Glowinski and Venance, 2005; Shen *et al*., 2008). Although, LTD is often considered the preponderant form of plasticity in the striatum (Paolo Calabresi, Maj, Pisani, *et al*., 1992; Kreitzer and Malenka, 2008; Wu *et al*., 2015), the direction of plasticity depends strongly on induction protocol and cellular context. In extracellular recordings without paired depolarisation, HFS principally triggers striatal LTD (Paolo Calabresi, Maj, Pisani, *et al*., 1992), whereas coincident postsynaptic depolarisation that relieves NMDAR Mg²⁺ block favours LTP (P. Calabresi *et al*., 1992). White matter stimulation has been shown to preferentially drive striatal LTP expression under such paired condition (Spencer and Murphy, 2000). In addition, LTP is more readily triggered in *in vivo* conditions (Charpier and Deniau, 1997). Thus, the induction protocol and local microenvironment critically determine the direction of plasticity expressed by SPNs.

Under our whole-cell paired HFS protocol, both LTP and LTD were observed in WT neurons, reflecting bidirectional plasticity. In contrast, eIF4E-TG SPNs exhibited a marked bias toward LTP, with an increased proportion of neurons expressing potentiation and an enhanced magnitude of LTP. These findings indicate that eIF4E overexpression shifts plasticity rules toward potentiation under these recording conditions.

Several mechanisms may contribute to this shift. The increased number of dendritic spines and elevated mEPSC frequency suggest a higher density of excitatory synapses, which may lower the threshold for potentiation. In addition, the reduction in AMPAR-mediated mEPSC amplitude raises the possibility that synapses may have reduced baseline AMPAR content, potentially allowing greater dynamic range for AMPAR insertion during LTP. The unchanged NMDA-mEPSC amplitude suggests that NMDAR-mediated currents are preserved, potentially altering the relative contribution of NMDAR versus AMPAR signalling during plasticity induction.

Importantly, blockade of NMDARs abolished LTP in both genotypes, indicating that this form of potentiation remains NMDAR-dependent. Thus, the enhanced LTP observed in eIF4E-TG SPNs does not represent a novel induction mechanism, but rather a shift in the probability and magnitude of NMDAR-dependent potentiation.

### Dopamine dependence and NMDAR gating of plasticity

Although dopamine release is reduced in the eIF4E-TG mice (Carbonell-Roig *et al*., 2024), it is still observed during HFS stimulation. The persistence of LTP in the presence of D1R and D2R antagonists therefore suggests that potentiation was not prevented by dopamine receptor blockade. This may reflect two possibilities: 1) that the LTP is not reliant on dopamine and thus is unaffected even in presence of blockers, or 2) that the D1- and D2-receptors signalling are altered in eIF4E-TG mice.

Interestingly, it was found that knockout of D2-receptors can reveal an HFS-induced, NMDAR-dependent LTP in striatal slices that is insensitive to D1R/D2R blockade (Calabresi *et al*., 1997). In WT mice, the same HFS protocol induces LTD and only induces LTP in Mg2+-free ACSF, and both directions are modifiable by D1/D2R antagonism (Calabresi *et al*., 1997). The LTP phenotype in D2R-knockout mice may be driven by homeostatic alterations in NMDAR and glutamatergic signalling and potentially reflects the role of D2R activation in inhibition of Ca^2+^ currents (Surmeier, Song and Yan, 1996; Calabresi *et al*., 1997), which are important for LTP expression in the striatum (Calabresi *et al*., 1996). It is therefore possible that altered dopamine receptor signalling may shift the balance of Ca²⁺-dependent plasticity mechanisms. However, because enhanced LTP was observed in both D1- and D2-SPNs, a mechanism affecting NMDAR-mediated signalling more broadly may be more consistent with our data.

The role of dopamine in striatal LTP has also been studied in parkinsonian mouse models. Although parkinsonian models such as Parkin and PINK1 knockout mice exhibit impaired HFS-induced LTP or LTD in the context of reduced striatal dopamine release (Kitada *et al*., 2007, 2009), eIF4E-TG mice display enhanced striatal plasticity (Santini et al., 2013) despite reduced dopamine signalling (Carbonell-Roig *et al*., 2024). These opposing phenotypes may reflect differences in the underlying molecular perturbation. Parkin and PINK1 regulate protein turnover pathways, including components of the ubiquitin–proteasome system, autophagy and mitophagy (Shimura *et al*., 2000; Gómez-Sánchez *et al*., 2016), whereas eIF4E overexpression selectively enhances cap-dependent translation initiation (Sonenberg and Gingras, 1998). Because eIF4E preferentially enhances translation of specific mRNA subsets, including transcripts involved in synaptic structure and function, altered translational selectivity, rather than generalised protein accumulation, may reshape the synaptic proteome of SPNs. Thus, differential regulation of protein synthesis and degradation may distinctly influence striatal plasticity outcomes. The balance between translational control and protein turnover may therefore be critical for maintaining normal plasticity in the striatum.

Although protein synthesis has been extensively linked to synaptic plasticity in hippocampal circuits, its contribution to striatal plasticity remains comparatively less characterised (Kelly, Mullany and Lynch, 2000; Banko *et al*., 2005; Gkogkas *et al*., 2013). We previously demonstrated increased rates of protein synthesis in the striatum of eIF4E-TG mice using the SUnSET assay (Aaltonen *et al*., 2026). While the present study does not directly test translation dependence during LTP induction or maintenance, the enhanced probability and magnitude of LTP observed here are consistent with a shift in plasticity thresholds that may arise from altered translational control. Such changes may facilitate stabilisation of potentiated synapses once induced, thereby biasing SPNs toward persistent potentiation. These findings extend the framework of translational regulation into striatal plasticity phenotypes relevant to neurodevelopmental disorders.

Striatal plasticity requires coordinated NMDAR-mediated Ca^2+^ influx (Calabresi *et al*., 1996; Shindou, Ochi-Shindou and Wickens, 2011; Jędrzejewska-Szmek *et al*., 2017) together with dopamine-dependent signalling (Paolo Calabresi, Maj, Mercuri, *et al*., 1992; Centonze *et al*., 1999; Reynolds and Wickens, 2002; Kreitzer and Malenka, 2005; Pawlak and Kerr, 2008; Shen *et al*., 2008; Reynolds *et al*., 2022). In our recordings, postsynaptic depolarisation during HFS would relieve the voltage-dependent Mg²⁺ block of NMDARs, promoting Ca²⁺ entry and favouring LTP induction (P. Calabresi *et al*., 1992; Fino *et al*., 2010). Although WT SPNs expressed bidirectional plasticity under this protocol, eIF4E-TG SPNs showed a bias toward potentiation that remained strictly NMDAR-dependent.

Given the established role of NMDAR-mediated dendritic Ca²⁺ microdomains in regulating synaptic plasticity (Carter and Sabatini, 2004), we examined whether Ca²⁺ signalling was altered in TG neurons. While absolute depolarisation-evoked Ca²⁺ signals were reduced in TG SPNs, the relative dendrite-to-soma Ca²⁺ ratio was increased, indicating altered compartmental distribution of Ca²⁺ signals. Because dendrites represent the principal site of synaptic integration and LTP expression, such redistribution may lower the threshold for potentiation without requiring an overall increase in global Ca²⁺ levels. Notably, NMDAR activation can under certain conditions drive sustained Ca²⁺ elevations that support dopamine-independent LTP (Park, Popescu and Poo, 2014), suggesting that sufficient postsynaptic Ca²⁺ signalling may overcome canonical dopamine dependence.

Translational dysregulation has been shown to impact NMDAR function in other ASD-related models, such as FXS mice, where altered NMDAR currents are associated with disrupted striatal LTP (Neuhofer *et al*., 2015). Together, these findings raise the possibility that altered dendritic Ca²⁺ integration in eIF4E-TG SPNs contributes to the observed bias toward NMDAR-dependent, dopamine-insensitive potentiation. Future studies will be required to determine whether exaggerated LTD in this model (Santini *et al*., 2013) similarly exhibits altered dopamine dependence and how compartmental Ca²⁺ dynamics interact with translational control to shape plasticity direction.

### Possible behavioural consequences of altered plasticity

As the induction protocol in our study generates LTP and LTD in WT neurons in roughly equal magnitudes and proportions, the average post-stimulation EPSC observed across neurons remained near baseline. This suggests that under our recording conditions, bidirectional plasticity in WT SPNs does not produce a net population-level shift in synaptic strength. In contrast, the bias toward potentiation observed in eIF4E-TG SPNs results in a net increase in excitatory drive at the population level. How such a shift translates to circuit and behavioural outcomes remains to be determined.

Corticostriatal plasticity is strongly linked to striatal-dependent learning processes (Di Filippo *et al*., 2009). Striatal LTP is associated with both procedural learning and goal-directed learning (Balleine, Delgado and Hikosaka, 2007; Yin et al., 2009; Hawes et al., 2013; Shan et al., 2014; Zhou et al., 2019), and synchronised SPN activity is particularly important for flexibility of goal-directed learning (Matamales *et al*., 2020; Peak *et al*., 2020). A persistent bias toward potentiation, particularly under conditions of reduced dopamine sensitivity, may alter updating of learned associations when contingencies change (Sala-Bayo *et al*., 2020; Fleps *et al*., 2025), resulting in more rigid behavioural outcomes. Such a mechanism is consistent with previously reported impairments in reversal learning in eIF4E-TG mice (Santini *et al*., 2013; Carbonell-Roig *et al*., 2024) and with broader evidence linking dopaminergic dysfunction to behavioural inflexibility in ASD models (Kosillo *et al*., 2019). However, direct causal links between the plasticity phenotype described here and behavioural outcomes remain to be established.

In conclusion, we identify an altered profile of striatal synaptic plasticity in eIF4E-TG mice characterised by increased spine density, enhanced spontaneous excitatory transmission, and a bias toward NMDAR-dependent potentiation. This potentiation was not prevented by D1/D2 receptor antagonism at the concentrations tested and was accompanied by altered compartmental Ca²⁺ dynamics. Together, these findings indicate that dysregulated translational control is associated with a shift in striatal plasticity rules. Such changes may contribute to altered circuit function and behavioural inflexibility observed in this model and highlight the importance of mRNA translational regulation in shaping striatal synaptic integration.

## Limitations of the study

The present study identifies an altered NMDAR-dependent plasticity profile in striatal projection neurons of eIF4E-TG mice; however, several limitations should be considered. First, although eIF4E overexpression increases translational capacity, protein synthesis was not directly manipulated during LTP induction or maintenance, and causality between altered translation and enhanced potentiation remains to be established. Second, plasticity experiments were conducted under disinhibited conditions with the addition of picrotoxin, and with single-concentration dopaminergic antagonism, limiting conclusions regarding dopamine dependence across physiological states. While the bath application of picrotoxin ensures minimal interference from inhibitory networks and removes the parameter of altered inhibitory connectivity in eIF4E-TG mice, this reduces the relevance to in vivo intact physiological conditions. Third, while white-matter stimulation recruits heterogeneous glutamatergic afferents, precluding pathway-specific attribution of the observed LTP. Addressing these questions will require input-selective induction protocols, translation-targeted manipulations, and experiments under intact inhibitory control.

## Methods

### Mice

All experiments were compliant with the ethical permits issued by the Swedish Board of Agriculture (Ethical number: 18194-2018 and 19345-2023) and performed in accordance with the European Parliament and Council Directive 210&63/EU, 22nd September 2010 for protection of animals used for scientific purposes. Mice were sex-separated and group-housed (up to five mice per cage) in a temperature (23 °C) and humidity (55 %) controlled environment, on a 12-hour light/dark cycle with water and food available *ad libitum*. In all cases, eIF4E^wt/wt^ mice are referred to as wild type (WT), and eIF4E^wt/βtEif4e^ mice are referred to as eIF4E-transgenic (eIF4E-TG or TG) (Ruggero *et al*., 2004; Santini *et al*., 2013). We used postnatal day 60-90 (2-3-month-old) male and female double-transgenic mice generated by crossing Drd1a-TdTomato^+/-^ (D1-TdTomato) mice (Shuen *et al*., 2008) (Jackson Laboratory, #016204) with eIF4E^wt/βtEif4e^ producing D1-TdTomato/WT and D1-TdTomato/TG, or by crossing Tg(Drd2-EGFP)S118Gsat^+/-^ (D2-eGFP) mice (Gong *et al*., 2003) with eIF4E^wt/βtEif4e^ producing D2-eGFP/WT and D2-eGFP/TG mice. Genotypes were confirmed by PCR as previously described (Carbonell-Roig *et al*., 2024; Aaltonen *et al*., 2026). All lines were maintained in hemizygosity by crossing with C57BL/6J mice purchased from Janvier. Due to the absence of differences between the identified D1- and D2-SPNs in all parameters, results from both SPN types have been combined. In addition, no differences were identified between males and females, so data have been combined.

### Stereotaxic surgery

Mice were injected with buprenorphine (i.p. 0.1 mg/kg) and anaesthetised with 2 % isoflurane. Adeno-associated virus (AAV) injections were performed via a Quintessential Stereotaxic Injector (Stoelting) at a rate of 30 nl/min. To assess the corticostriatal input strength, bilateral viral injections of 200 nl pAAV-hSyn-hChR2(H134R)-mCherry (Addgene #26976) were administered in the primary motor cortex (AP +1.5, ML +/-1.8, DL -0.7) or somatosensory cortex (AP -1.5, ML +/-3.0, DL -0.7) of WT and eIF4E-TG mice. For the Ca^2+^ imaging experiment, 250 nl pGP-AAV-syn-FLEX-jGCaMP7s-WPRE (Addgene #104491) was bilaterally injected in the striatum (AP +0.8, ML - 1.8, DV -2.4). Post-operative carprofen and buprenorphine were administered i.p. twice a day for 24 hours (5 mg/kg and 0.1 mg/kg, respectively). Mice were allowed to recover for at least three weeks to permit sufficient viral expression.

### Electrophysiology

#### Sample preparation

Acute brain slices were prepared using either sucrose (in mM: 10 NaCl, 2.5 KCl, 25 NaHCO3, 1.25 NaH2PO4, 7 MgCl2, 0.5 CaCl2, 180 sucrose, 10 glucose) for mEPSC/mIPSC recordings and fast-scan cyclic voltammetry (FSCV), or NMDG-based cutting solutions (Ting *et al*., 2018) (in mM: 92 NMDG, 30 NaHCO3, 2.5 KCl, 20 HEPES, 2 thiourea, 1.25 NaH2PO4, 3 Na-pyruvate, 5 ascorbic acid, 25 glucose, 0.5 CaCl2, 10 MgCl2) titrated with HCl to pH 7.4, for NMDA-mEPSC, optogenetic and plasticity recordings. The slices were then stored in standard ACSF (in mM: 125 NaCl, 2.5 KCl, 25 NaHCO3, 1.25 NaH2PO4, 1 MgCl2, 2 CaCl2, 10 glucose) or HEPES-based holding solution, respectively (in mM: 92 NaCl, 30 NaHCO3, 2.5 KCl, 20 HEPES, 2 thiourea, 1.25 NaH2PO4, 3 Na-pyruvate, 5 ascorbic acid, 25 glucose, 2.5 CaCl2, 2 MgCl2 titrated with NaOH to pH 7.4). Coronal sections (250 µm) containing the striatum were prepared using a Leica vibratome (VT 1200S). Sucrose-prepared slices were incubated at 32-34 °C for 30 minutes in oxygenated ACSF, whereas NMDG-prepared slices were stored in NMDG solution for 10 minutes at 32-34 °C before transferring to HEPES holding solution. Slices were transferred to a recording chamber, where they were continuously perfused with oxygenated ACSF at a rate of 3 ml/min.

#### Recording protocol

Recordings were obtained using a patch clamp electrophysiology rig fitted with SciCam pro camera (Scientifica, United Kingdom) equipped with a 40 × 0.8 NA water-immersion objective (LUMPlanFLN, Olympus, United States) and Dodt contrast tube optics. Cell/slice visualisation and light delivery were performed using a CoolLED PE-300 Ultra LED. Recordings were obtained with a Multiclamp 700B amplifier (Molecular Devices) and Axon Digidata 1550B digitizer (Molecular Devices, United States), using pCLAMP 11 software (Molecular Devices, United States). Data were low pass filtered at 2-10 kHz depending on protocol. For all recordings, borosilicate glass capillaries were used to prepare electrodes with 3–5 MΩ tip resistance. Recordings were not corrected for liquid junction potentials, and series resistance compensation was not applied during any of the recordings. For all recordings, series resistance was monitored and recordings were excluded if series resistance exceeded 30 MΩ or changed by over 20%, or in the event of declining cell health.

All recordings were done at 30-34 °C. To detect miniature glutamate-mediated excitatory synaptic currents (mEPSCs), electrodes were filled with intracellular solution (IS) containing the following (in mM): 115 K-gluconate, 20 KCl, 20 HEPES, 1 MgCl2, 2 MgATP, 0.2 NaGTP, adjusted to pH 7.25 with KOH. To detect miniature GABA-mediated inhibitory synaptic currents (mIPSCs), 20 mM K-gluconate plus 115 mM KCl was substituted for 115 mM K-gluconate and 20 KCl using the previous IS. All other recordings were obtained using a caesium methanesulfonate-based internal solution containing (in mM): 126 CsMeSO3, 20 KCl, 4 MgATP, 0.3 Na2-GTP, 8 Na2-phosphocreatine, 1 EGTA, 10 HEPES and 0.1 CaCl2 at pH 7.25 and osmolarity 300 mOsm.

Recordings of AMPA-receptor derived mEPSCs were done with bath application of TTX (1 µM) to block action potential formation and propagation, and picrotoxin (100 µM) was added to IS to block postsynaptic GABAergic receptors. Recordings of mIPSCs were performed with bath application of TTX (1 µM), and DNQX (10 µM) and DL-AP5 (40 µM) were included to block glutamatergic currents. NMDA-receptor derived mEPSCs were recorded with bath application of TTX (300 nM), picrotoxin (50 µM) and DNQX (10 µM) in Mg^2+^-free ACSF. Optogenetic recordings, plasticity and 2-photon recordings were done in the presence of picrotoxin (50 µM). D-AP5 (10 µM), SCH39166 (1 µM) and sulpiride (2 µM) were added to the recording solution in a subset of plasticity recordings to block NMDA-receptors, D1 or D2 dopamine receptors, respectively. Chemicals were purchased from Fisher Scientific, Tocris and HelloBio.

Whole-cell voltage-clamp recordings were performed on striatal neurons identified by TdTomato or eGFP fluorescence. For all recordings, the access resistance was monitored throughout the recording by applying hyperpolarizing 10 mV voltage steps. Recordings were discarded if access resistance exceeded 30 MΩ or changed by over 20 %. All recordings were performed in voltage-clamp configuration, with membrane potential clamped at -70 mV unless stated otherwise and were not corrected for liquid junction potential. Miniature excitatory and inhibitory post-synaptic currents (EPSC and IPSC) recordings were obtained in gap-free mode for five-10 minutes. Optogenetic recordings were obtained using either single or paired (delivered 100 ms apart) 1 ms pulses of blue light from a CoolLED PE-300 Ultra LED module through the 40x objective, at a rate of 0.033 Hz. Light intensity was set for each cell to provide an average EPSC of 200 pA. For NMDA: AMPA recordings, light stimulation was first given at -70 mV holding potential for several sweeps, before raising the holding potential to +40 mV and recording for several more sweeps.

Plasticity recordings were performed using paired (50 ms) electrical stimulation at a rate of 0.05 Hz. Baseline EPSC was established over 10 minutes, and stimulation intensity was set for each cell to provide an average EPSC of 250 pA. Cells exhibiting unstable baseline were excluded from recordings. After a stable baseline recording was obtained, plasticity induction stimulation was given in the form of high-frequency stimulation at 100 Hz for one second, repeated four times with 10-second intervals. Each one-second bout of high-frequency stimulation (HFS) was paired with postsynaptic depolarisation of the holding potential to 0 mV. The stimulation was increased to twice the intensity used during baseline stimulation and reduced to the original level for recording the post-stimulation response.

#### Fast-scan cyclic voltammetry

Carbon fibre electrodes were generated as previously described (Carbonell-Roig *et al*., 2024) and filled with 1 M KCl. Electrodes were calibrated on each day of recording by measuring the response to known concentrations of dopamine (0.5, 1 and 2 µM). Striatal slices were placed in the recording chamber and supplied with a constant flow of oxygenated (95 % O2/ 5 % CO2) ACSF at 3 ml/min, maintained at 30-34 °C.

Recordings were obtained using the Chem-clamp 5MEG amplifier (Dagan corporation, Minnesota, USA), using the Demon Voltammetry suite software (Yorgason, España and Jones, 2011). A triangular voltage wave (−0.4V to +1.2V at 400V/s) was applied to the slice through the recording electrode at a frequency of 0.1 Hz. Slices were stimulated with a bipolar electrode placed 100-200 μm from the recording electrode, using an Iso-Flex stimulus isolator (A.M.P.I.). Single pulses (1 ms) were applied every 90 seconds to determine baseline dopamine release. As above, HFS was applied at a frequency of 100 Hz for one second, repeated four times with 10-second intervals. Slices were excluded if baseline or post-HFS dopamine currents were unstable or below detection threshold.

#### Two-photon GCaMP7s-based calcium imaging

Calcium imaging experiments were performed on a Scientifica Hyperscope multiphoton imaging system (Scientifica, Uckfield, UK) equipped with a 16x water-immersion objective (Zeiss, NA 0.8). Two-photon excitation of GCaMP7s was provided by a Chameleon Vision 1 femtosecond Ti-Sapphire laser (Coherent) tuned to 920 nm. Emitted fluorescence was collected through ET525/50m emission filters (Chroma Technologies) and detected by bi-alkali photomultiplier tubes (PMTs; R9880U, Hamamatsu). Laser power at the sample plane was controlled via a voltage-controlled Pockels cell and maintained below 21 mW to minimise phototoxicity and photobleaching. Images were acquired using a resonant scanner in ScanImage software (version 2021.0.0.3baa4b74be) at a frame rate of 5.3 Hz with a zoom factor of 15, resulting in 5.04 pixels per µm.

To evoke somatic depolarisation, we gained voltage control using whole-cell patch clamp recordings of neurons visualised using infrared Dodt gradient contrast imaging and an AxioCam ICm1 camera (Zeiss) controlled with ZEN 3.4 software. Neurons were impaled with patch pipettes and filled with caesium methanosulfonate-based internal solution (see electrophysiology > recording protocol). SeTau-647 (30 μM, Hölzel biotech) was included in the pipette to fluorescently mark the recorded neuron. Neurons were voltage-clamped at −70 mV and intermittent step commands (500 ms) depolarised the neuron to 0 mV. Step commands and membrane voltage recordings were performed with WinWCP (StratChlyde University, v5.8.3) controlling an Axopatch 200B amplifier (Molecular Devices) via a vDAQ data acquisition interface (National Instruments). During depolarisation, a trigger signal was sent to ScanImage (MATLAB-based software) to synchronise the onset of two-photon image acquisition with the electrophysiological stimulation.

#### Analysis

All analysis of patch-clamp data was performed using Clampfit or the open-access Python-based software Stimfit (Guzman, Schlögl and Schmidt-Hieber, 2014), whereas FSCV data analysis was performed using Demon Voltammetry suite software (Yorgason, España and Jones, 2011). Miniature EPSCs and IPSC recordings were analysed using template scaling for event detection (Clements and Bekkers, 1997). Events were filtered using a post-analysis threshold of four times the standard deviation of the baseline of each detected event. Evoked responses were analysed from averages of at least 10 sweeps. Paired pulse ratio was analysed by expressing the amplitude of the second EPSC over the first EPSC. NMDA:AMPA ratio was analysed by expressing the amplitude of the +40 mV EPSC, 50 ms after stimulation, divided by the -70 mV EPSC. In plasticity recordings, the post-stimulation EPSC (first EPSC from the paired response) was calculated as an average of the EPSCs at minutes 25-30 after the plasticity induction protocol, which was then expressed as a percentage change from the average baseline EPSC (100 %) during 10 minutes prior to stimulation. If the EPSC changed from baseline by over 10 % (increase or decrease) at 30 minutes post-stimulation, the cell was considered to express long-term plasticity.

FSCV recordings of HFS-evoked dopamine release were analysed by expressing the size of the first peak elicited with HFS as a percentage change from the average baseline dopamine release (100 %) elicited from five sweeps prior to HFS.

Two-photon GCaMP7s image analysis was performed using ImageJ software (v1.54p). Soma and dendrites were defined as regions of interest (ROIs), and mean fluorescence intensity values were extracted from each ROI across the time series. Background fluorescence was subtracted from each frame prior to further analysis. Relative changes in fluorescence (ΔF/F₀) were calculated over time. Dendritic ROIs were selected at distances greater than 60 µm from the soma, with a maximum of five dendritic measurements per neuron.

### Immunofluorescence

The process of microinjection was followed as previously described (Dumitriu, Rodriguez and Morrison, 2011; Aaltonen et al., 2025). Mice underwent cardiac perfusion with 4 % paraformaldehyde (PFA) in 0.1 M PBS (pH 7.4), and brains were fixed for a further four hours at 4 °C, before slicing at 200 µm using a Leica VT100s vibratome. Striatal cells were filled with 8 % Lucifer Yellow CH lithium salt (Invitrogen) in PBS and Tris-HCl via current injection of 2-5 nA over 5-10 minutes. To amplify the fluorescent signal from biotin-conjugated Lucifer Yellow, slices were incubated for 48 hours in Streptavidin Alexa Fluor™-488 (Invitrogen, 1:200) in 0.6 % Triton X-100 in TBS. Slices were incubated overnight in 2 % NGS / 0.6 % TBS-triton with goat anti-mouse Alexa Fluor™ 488 (Invitrogen, 1:500). Dendritic sections were imaged with a Zeiss LSM 800-Airy system microscope with a 63x /1.4 oil immersion objective (Art.Nr 420782-9900-799). Dendritic spine analysis was performed using the Imaris software 9.5.1 (Oxford Instruments).

To assess the specificity of channelrhodopsin expression in the optogenetic electrophysiological experiment, 30-40 µm coronal brain slices were prepared in PBS. Slices containing striatum were selected for immunostaining. Slices were permeabilised for five minutes in TBS containing 1:10 methanol and 1:3 hydrogen peroxide. A second permeabilisation step was performed using TBS with 0.2 % Triton X-100 (TBS-Triton) and incubated for 15 minutes. Slices were incubated with primary antibody overnight at 4 °C (Rockland 600-401-379 rabbit anti-RFP; 1:1000) and then incubated with secondary antibody for 45-60 minutes at room temperature (A-11011 anti-rabbit 568; 1:500). Sections were imaged at 10x with a Zeiss LSM800 confocal microscope, and images were analysed using the DMC-BrainMAP Napari plug-in (Jung *et al*., 2025).

### Statistical analysis

All statistical analyses were performed using GraphPad Prism. N-values indicate number of mice and n-values indicate the number of cells or slices, and data are presented as median ± quartiles or mean ± SEM. Data distribution was assessed for normality prior to parametric testing. Two-tailed Student’s t-tests or Mann-Whitney U-tests were used for two-group comparisons, whereas two-way ANOVAs were used for grouped variable comparisons, with Tukey’s multiple comparisons tests performed for post-hoc comparisons between groups. Multiple comparisons tests were performed only in the case of an interaction effect. Chi-squared tests were used to compare qualitative variables between genotypes. In all cases, * p<0.05, ** p<0.01, *** p<0.001 and **** p<0.0001, ns p>0.05. Interaction effects were denoted as # p<0.05, ## p<0.01, ### p<0.001 and #### p<0.0001. Key statistical parameters, including N and n values, mean ± SEM, median ± quartiles, t-, u- and f-values, p-values and significance levels are indicated for each statistical test in Supplementary tables 1-6.

## Acknowledgements

We thank the past and current members of Borgkvist and Santini labs for their methodological support and regular insightful discussions. We would like to thank Gilad Silberberg, Maya Ketzef and other members of the neuroscience department for their considerate feedback and advice. We are grateful to Gilberto Fisone for providing the Tg(Drd2-EGFP)S118Gsat^+/-^ mice. We extend further thanks to the staff and veterinarians of the Comparative Medicine Biomedicum (KM-B) for their continuous assistance in maintaining the mouse colonies.

## Author contributions

Conceptualisation, A.B., E.S. and E.C.; Methodology, A.A., A.B. and E.S.; Validation, A.A., A.B. and E.S.; Formal Analysis, A.A., A.B. and E.S.; Investigation, A.A., J.R.L., J.O., C.C., O.L., A.B. and E.S.; Resources, A.B. and E.S.; Writing – Original Draft, A.A., C.C., J.O., A.B. and E.S.; Writing – Review & Editing, A.A., A.B. and E.S.; Visualisation, A.A, A.B. and E.S.; Supervision, A.B. and E.S.; Project Administration, A.B. and E.S.; Funding Acquisition, A.B., E.S., E.C.

## Supplementary figures

**Supplementary figure 1.**
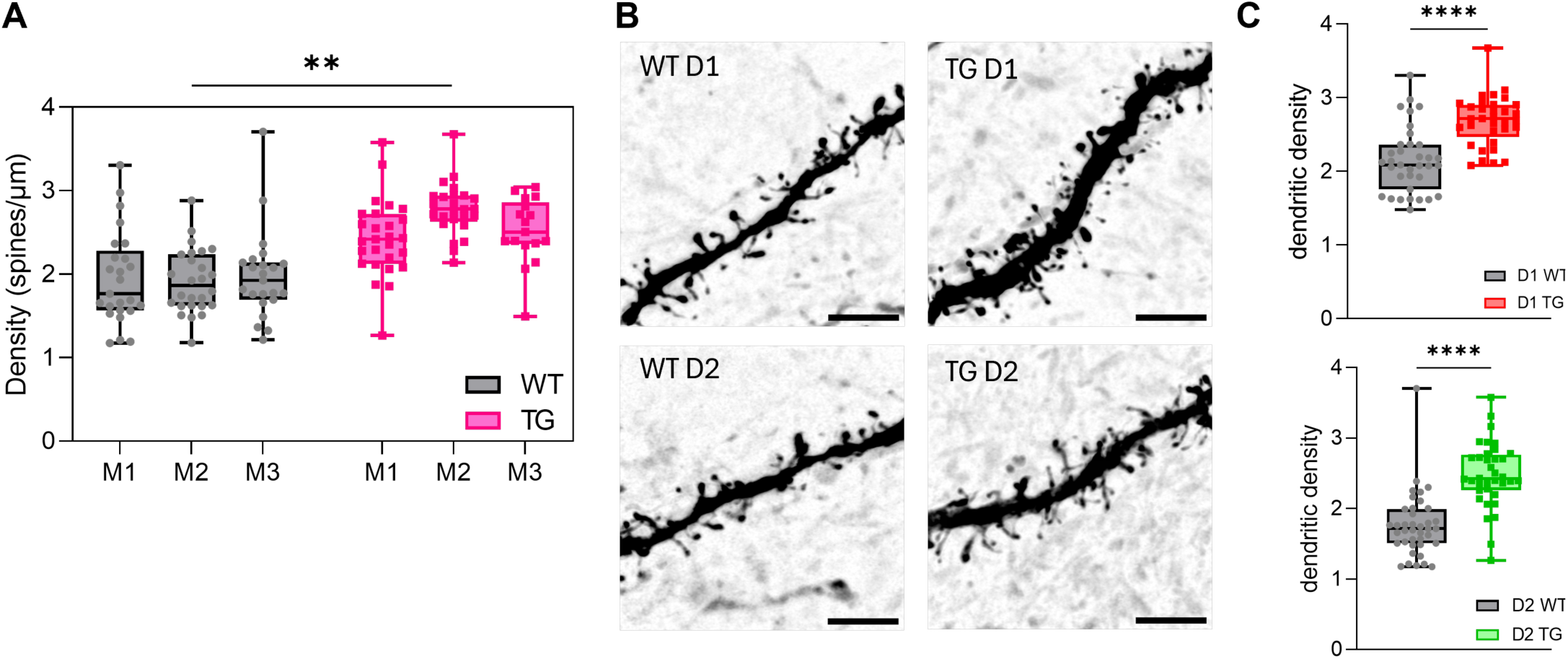
Dendritic spines of D1- and D2-SPNs from eIF4E-TG mice. **A)** Nested analysis of the results shown in Figure 1 (nested T-test: t=5.818, **p<0.0043). Each dot represents a dendrite, and each box plot represents a mouse. **B-C)** Images were collected from 3 WT and 3 TG mice (D1: dendrites n=35 WT, n=33 TG; D2: dendrites n=39 WT, n=36 TG). **A)** Example Airyscan confocal images of D1- and D2-SPN dendrites (scale bar 4µm). **B)** D1-SPN dendritic spine density in eIF4E-TG mice (unpaired T-test: t=5.521, ****p<0.0001). **C)** D2-SPN dendritic spine density in eIF4E-TG mice (unpaired T-test: t=6.835, ****p<0.0001). Data are presented as median ± quartiles within the box plot, with whiskers indicating minimum and maximum values. Dots represent individual values for each dendrite. Significance is denoted as ** p<0.005, **** p<0.0001, calculated using Student’s or nested t-test. For full details of statistical analysis, including negative results, refer to Supplementary table 6.

**Supplementary table 1.**
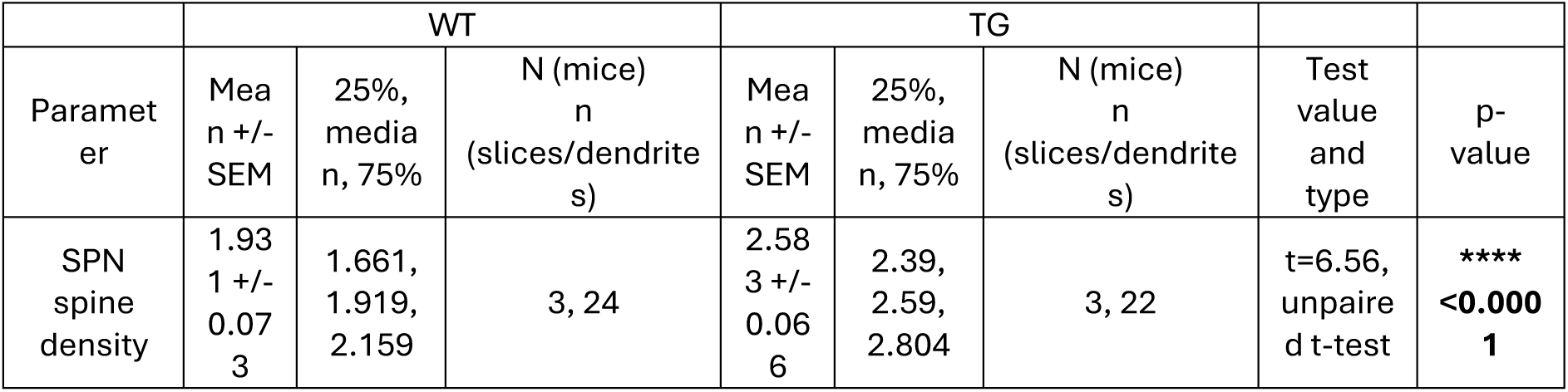

**Supplementary table 2.**

|  | WT |  |  | TG |  |  |  |  |
| --- | --- | --- | --- | --- | --- | --- | --- | --- |
| Parameter | Mean<br>+/-<br>SEM | 25%,<br>median,<br>75% | N<br>(mice)<br>n<br>(cells) | Mean<br>+/-<br>SEM | 25%,<br>median,<br>75% | N<br>(mice)<br>n<br>(cells) | Test value<br>and type | p-value |
| mEPSC<br>frequency | 1.025<br>+/-<br>0.117 | 0.722,<br>0.906,<br>1.356 | 7, 19 | 2.336<br>+/-<br>0.21 | 1.822,<br>2.390,<br>2.951 | 5, 16 | t=5.665,<br>unpaired t-test | ****<br>**<0.0001** |
| mEPSC<br>amplitude | 14.32<br>+/-<br>0.953 | 10.38,<br>14.45,<br>18.33 | 7, 19 | 10.68<br>+/-<br>0.693 | 8.527,<br>10.81,<br>12.78 | 5, 16 | t=2.985,<br>unpaired t-test | **<br>**0.0053** |
| mIPSC<br>frequency | 1.066<br>+/-<br>0.134 | 0.689,<br>0.99,<br>1.134 | 8, 17 | 1.044<br>+/-<br>0.102 | 0.742,<br>1.072,<br>1.165 | 5, 16 | t=0.129,<br>unpaired t-test | ns<br>0.8981 |
| mIPSC<br>amplitude | 32.03<br>+/-<br>1.57 | 27.78,<br>32.6,<br>36.5 | 8, 17 | 21.45<br>+/-<br>1.692 | 15.39,<br>21.33,<br>26.38 | 5, 15 | t=4.587,<br>unpaired t-test | ****<br>**<0.0001** |
| NMDA<br>mEPSC<br>frequency | 3.434<br>+/-<br>0.438 | 1.89,<br>3.07,<br>5.00 | 5, 17 | 5.473<br>+/-<br>0.526 | 3.55,<br>6.15,<br>7.18 | 6, 15 | t=3,<br>unpaired t-test | **<br>**0.0054** |
| NMDA<br>mEPSC<br>amplitude | 8.838<br>+/-<br>0.418 | 7.475,<br>8.67,<br>9.97 | 5, 17 | 9.883<br>+/-<br>0.59 | 8.81,<br>10.41,<br>10.52 | 6, 15 | t=1.47,<br>unpaired t-test | ns<br>0.1519 |

**Supplementary table 3.**
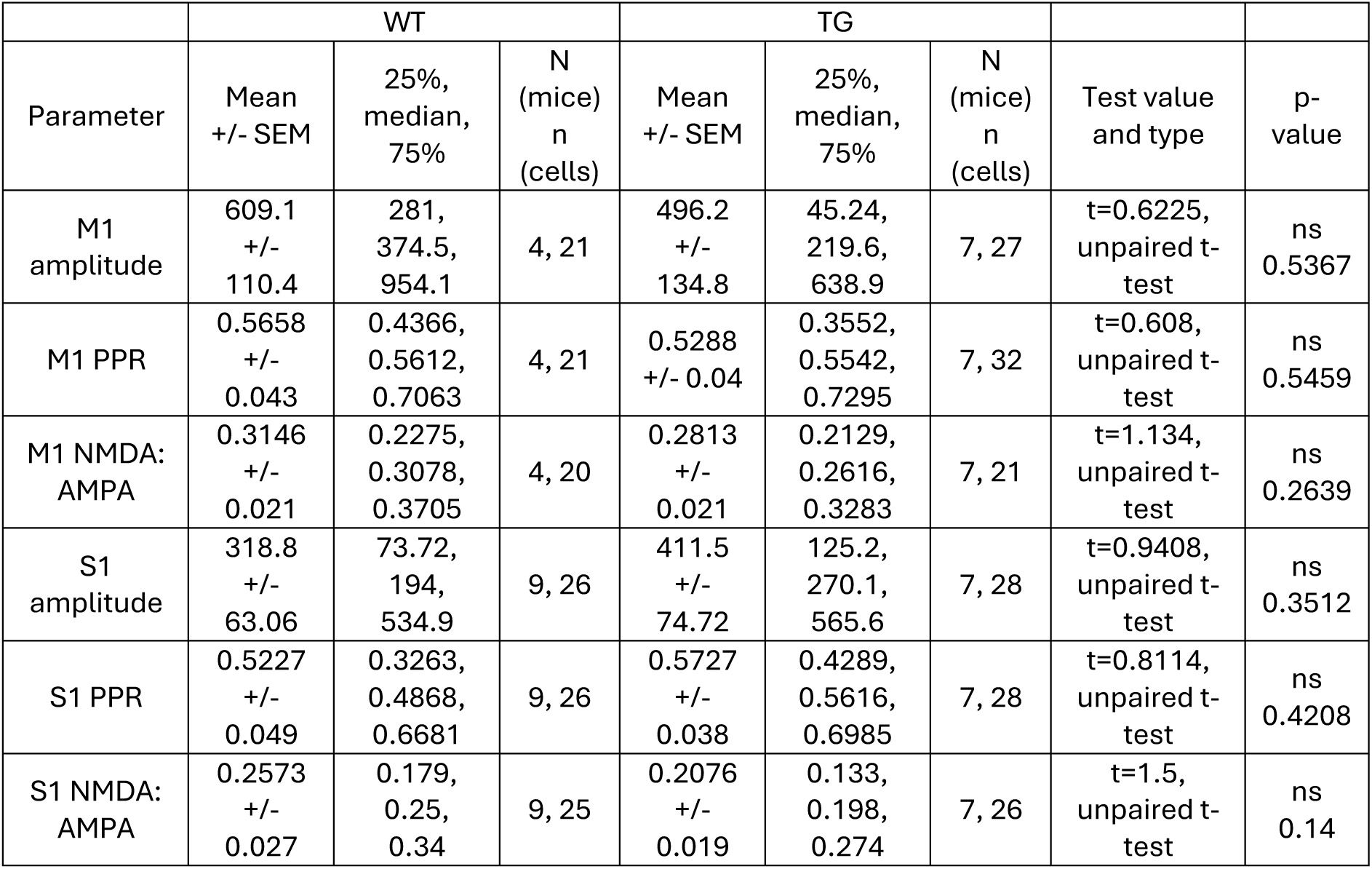

**Supplementary table 4.**

|  | WT |  |  | TG |  |  |  |  |
| --- | --- | --- | --- | --- | --- | --- | --- | --- |
| Parameter | Mean<br>+/- SEM | 25%,<br>median,<br>75% | N<br>(mice)<br>n<br>(cells) | Mean<br>+/- SEM | 25%,<br>median,<br>75% | N<br>(mice)<br>n<br>(cells) | Test value<br>and type | p-value |
| Plasticity<br>distribution | | | | | | | $\chi^2=4.413$,<br>Chi-<br>square<br>test | **\***<br>**0.0418** |
| EPSC change<br>(LTP) | 137.5<br>+/-<br>7.328 | 116.9,<br>135.7,<br>145.7 | 10, 11 | 235.1<br>+/-<br>28.79 | 148.9,<br>208.7,<br>276.3 | 12, 12 | t=3.157,<br>unpaired<br>t-test | **\*\***<br>**0.0048** |
| EPSC change,<br>ACSF | 107.395<br>+/-<br>9.017 | 83.713,<br>111.55,<br>135.77 | 21, 21 | 212.986<br>+/-<br>28.82 | 146.61,<br>189.67,<br>269.43 | 13, 14 | Two-way-ANOVA<br>Genotype:<br>$F_{(2, 65)}=20.60$;<br>Drug: $F_{(2, 65)}=1.5$;<br>Genotype<br>x Drug: $F_{(2, 65)}=3.738$ | Genotype:<br>**\*\*\*\***<br>**<0.0001**;<br>Drug: ns<br>0.2307;<br>Genotype<br>x Drug: **\***<br>**0.0291** |
| EPSC change,<br>AP5 | 124.077<br>+/-<br>13.03 | 96.044,<br>106.67,<br>160.61 | 7, 9 | 136.405<br>+/-<br>17.97 | 90.261,<br>128.28,<br>171.86 | 8, 11 |  |  |
| EPSC change,<br>SCH/sulpiride | 99.656<br>+/-<br>2.917 | 95.229,<br>97.067,<br>105.553 | 4, 8 | 194.129<br>+/-<br>16.032 | 154.613,<br>207.93,<br>227.033 | 4, 8 |  |  |
| Baseline DA<br>release | 0.6075<br>+/-<br>0.0556 | 0.4489,<br>0.5279,<br>1.014 | 4, 14 | 0.3753<br>+/-<br>0.0455 | 0.2533,<br>0.3604,<br>0.4347 | 4, 17 | t=3.262,<br>unpaired<br>t-test | **\*\***<br>**0.0028** |
| HFS DA<br>release % | 68.07<br>+/-<br>14.08 | 28.05,<br>55.25,<br>109.4 | 4, 14 | 60.63<br>+/-<br>9.024 | 39.3,<br>60.71,<br>90.16 | 4, 17 | t=0.4596,<br>unpaired<br>t-test | ns<br>0.6492 |

**Supplementary table 5.**

|  | WT |  |  | TG |  |  |  |  |
| --- | --- | --- | --- | --- | --- | --- | --- | --- |
| Parameter | Mean<br>+/- SEM | 25%,<br>median,<br>75% | N<br>(mice)<br>n<br>(cells) | Mean<br>+/- SEM | 25%,<br>median,<br>75% | N<br>(mice)<br>n<br>(cells) | Test<br>value and<br>type | p-value |
| Dendritic:<br>somatic<br>GCaMP7s peak<br>fluorescence | 1.302<br>+/-<br>0.2696 | 0.2616,<br>0.7021,<br>2.178 | 4, 30 | 2.274<br>+/-<br>0.2664 | 0.6892,<br>1.858,<br>3.166 | 5, 47 | u=422,<br>Mann-Whitney | **\*\***<br>**0.0051** |

**Supplementary table 6.**

|  | WT |  |  | TG |  |  |  |  |
| --- | --- | --- | --- | --- | --- | --- | --- | --- |
| Parameter | Mean<br>+/-<br>SEM | 25%,<br>median,<br>75% | N (mice)<br>n<br>(dendrites) | Mean<br>+/-<br>SEM | 25%,<br>median,<br>75% | N (mice)<br>n<br>(dendrites) | Test<br>value and<br>type | p-value |
| Nested<br>spine<br>density | 1.944<br>+/-<br>0.057 | 1.624,<br>1.871,<br>2.189 | 3, 74 | 2.579<br>+/-<br>0.05 | 2.35,<br>2.625,<br>2.842 | 3, 69 | t=5.818,<br>nested t-test | **\*\***<br>**0.0043** |
| D1-SPN<br>spine<br>density | 2.152<br>+/-<br>0.075 | 1.759,<br>2.09,<br>2.365 | 3, 35 | 2.684<br>+/-<br>0.059 | 2.469,<br>2.715,<br>2.906 | 3, 33 | t=5.521,<br>unpaired<br>t-test | ****<br>**<0.0001** |
| D2-SPN<br>spine<br>density | 1.758<br>+/-<br>0.073 | 1.507,<br>1.714,<br>1.991 | 3, 39 | 2.482<br>+/-<br>0.077 | 2.263,<br>2.426,<br>2.765 | 3, 36 | t=6.835,<br>unpaired<br>t-test | ****<br>**<0.0001** |

